# An engineered 4-1BBL fusion protein with “activity-on-demand”

**DOI:** 10.1101/2020.04.29.068171

**Authors:** Jacqueline Mock, Marco Stringhini, Alessandra Villa, Dario Neri

## Abstract

Engineered cytokines are gaining importance for cancer therapy but those products are often limited by toxicity, especially at early time points after intravenous administration. 4-1BB is a member of the tumor necrosis factor receptor superfamily, which has been considered as a target for therapeutic strategies with agonistic antibodies or using its cognate cytokine ligand, 4-1BBL. Here we describe the engineering of an antibody fusion protein (termed F8-4-1BBL), which does not exhibit cytokine activity in solution but regains biological activity upon antigen binding. F8-4-1BBL bound specifically to its cognate antigen, the alternatively-spliced EDA domain of fibronectin, and selectively localized to tumors *in vivo*, as evidenced by quantitative biodistribution experiments. The product promoted a potent anti-tumor activity in various mouse models of cancer, without apparent toxicity at the doses used. F8-4-1BBL represents a prototype for antibody-cytokine fusion proteins, which conditionally display “activity-on-demand” properties at the site of disease upon antigen binding and reduce toxicity to normal tissues.

## INTRODUCTION

Cytokines are immunomodulatory proteins, which have been considered for pharmaceutical applications for the treatment of cancer patients(1–3) and other types of disease(2). There is a growing interest in the use of engineered cytokine products as anti-cancer drugs, capable of boosting the action of T cells and natural killer (NK) cells against tumors(3, 4), alone or in combination with immune check-point inhibitors(3, 5–7).

Recombinant cytokine products on the market include IL2 (Proleukin^®^)(8, 9), IL11 (Neumega^®^)(10, 11), TNF (Beromun^®^)(12), IFNα (Roferon A^®^, Intron A^®^)(13, 14), IFNβ (Avonex^®^, Rebif^®^, Betaseron^®^)(15, 16) IFNγ (Actimmune^®^)(17), G-CSF (Neupogen^®^)(18), GM-CSF (Leukine^®^)(19, 20). The recommended dose is typically very low (often at less than one milligram per day)(21–23), as cytokines may exert biological activity in the subnanomolar concentration range(24). In order to develop cytokine products with improved therapeutic index, various strategies have been proposed. Protein PEGylation or Fc fusions may lead to prolonged circulation time in the bloodstream, allowing the administration of low doses of active payload(25, 26). In some implementation, cleavable PEG polymers may be considered, yielding prodrugs which regain activity at later time points(27). Alternatively, tumor-homing antibody fusions have been developed, since the preferential concentration of cytokine payloads at the tumor site has been shown in preclinical models to potentiate therapeutic activity, helping spare normal tissues(28–34). Various antibody-cytokine fusions are currently being investigated in clinical trials for the treatment of cancer and of chronic inflammatory conditions [for reviews, see(2, 33, 35–37)].

Antibody-cytokine fusions display biological activity immediately after injection to patients, which may lead to unwanted toxicity and prevent escalation to therapeutically active dose regimens(9, 22, 38). In the case of pro-inflammatory payloads (e.g., interleukin-2, interleukin-12, tumor necrosis alpha), common side effects include hypotension, nausea and vomiting, as well as flu-like symptoms(24, 39–42). These side-effects typically disappear when the cytokine concentration drops below a critical threshold, thus providing a rationale for slow-infusion administration procedures(43). It would be highly desirable to generate antibody-cytokine fusion proteins with excellent tumor targeting properties and with “activity-on-demand” (i.e., with a biological activity which is conditionally gained upon antigen binding at the site of disease, helping spare normal tissues).

Here, we describe a novel fusion protein, consisting of the F8 antibody (specific to the alternatively-spliced EDA domain of fibronectin(44, 45)) and of murine 4-1BBL, which did not exhibit cytokine activity in solution but could regain potent biological activity upon antigen binding. The antigen is conserved from mouse to man(46), is virtually undetectable in normal adult tissues (exception made for placenta, endometrium and some vessels in the ovaries), but is expressed in the majority of human malignancies(44, 45, 47, 48). 4-1BBL is a member of the tumor necrosis factor superfamily(49). It is expressed on antigen-presenting cells(50, 51) and binds to its receptor 4-1BB which is upregulated on activated cytotoxic T cells(52), activated dendritic cells(52), activated NK and NKT cells(53) and on regulatory T cells(54). Signaling through 4-1BB on cytotoxic T cells protects them from activation-induced cell death and skews the cell towards a more memory-like phenotype(55, 56).

We engineered nine formats of the F8-4-1BBL fusion protein and one of them exhibited a superior performance in quantitative biodistribution studies and conditional gain of cytokine activity upon antigen binding. The fusion protein was potently active against different types of cancer, without apparent toxicity at the doses used. F8-4-1BBL may represent a prototype for next-generation antibody-cytokine fusions with “activity-on-demand”. The EDA domain of fibronectin is a particularly attractive antigen for cancer therapy, in view of its high selectivity, stability and abundant expression in most tumor types(44, 45, 47, 48).

## RESULTS

Human 4-1BBL is a homotrimeric protein [**Figure 1a**](57), while its murine counterpart forms stable homodimers(58, 59). Stable trimeric structures can be engineered by connecting 4-1BBL monomeric domains with suitable polypeptide linkers(60). Recombinant antibodies can be expressed as full IgG or as fragments, forming single-chain Fv (scFv)(61, 62) or diabody(63) structures [**Figure 1b** and **Supplementary Table 1**](2, 64, 65). Nine different fusion proteins containing F8 antibody and murine 4-1BBL moieties were expressed in mammalian cells, in order to identify products with promising features for subsequent *in vivo* investigations. Mutational scans had revealed that the disulfide bond linking two 4-1BBL monomers is crucial for protein stability [**Supplementary Figure 1**]. The observation that the TNF homology domain (THD) within 4-1BBL was sufficient for full *in vitro* activity [**Supplementary Figure 2**] guided the design of the modules to be included in the fusion proteins. Six out of nine products exhibited favorable size exclusion and SDS-PAGE profiles [**Figure 1c** and **Supplementary Figure 3**]. We selected formats **2, 3, 5, 7** and **8** for further investigations, since those proteins gave the best yields and did not show signs of aggregation even after repeated freeze-thaw cycles.

**Figure 1:**
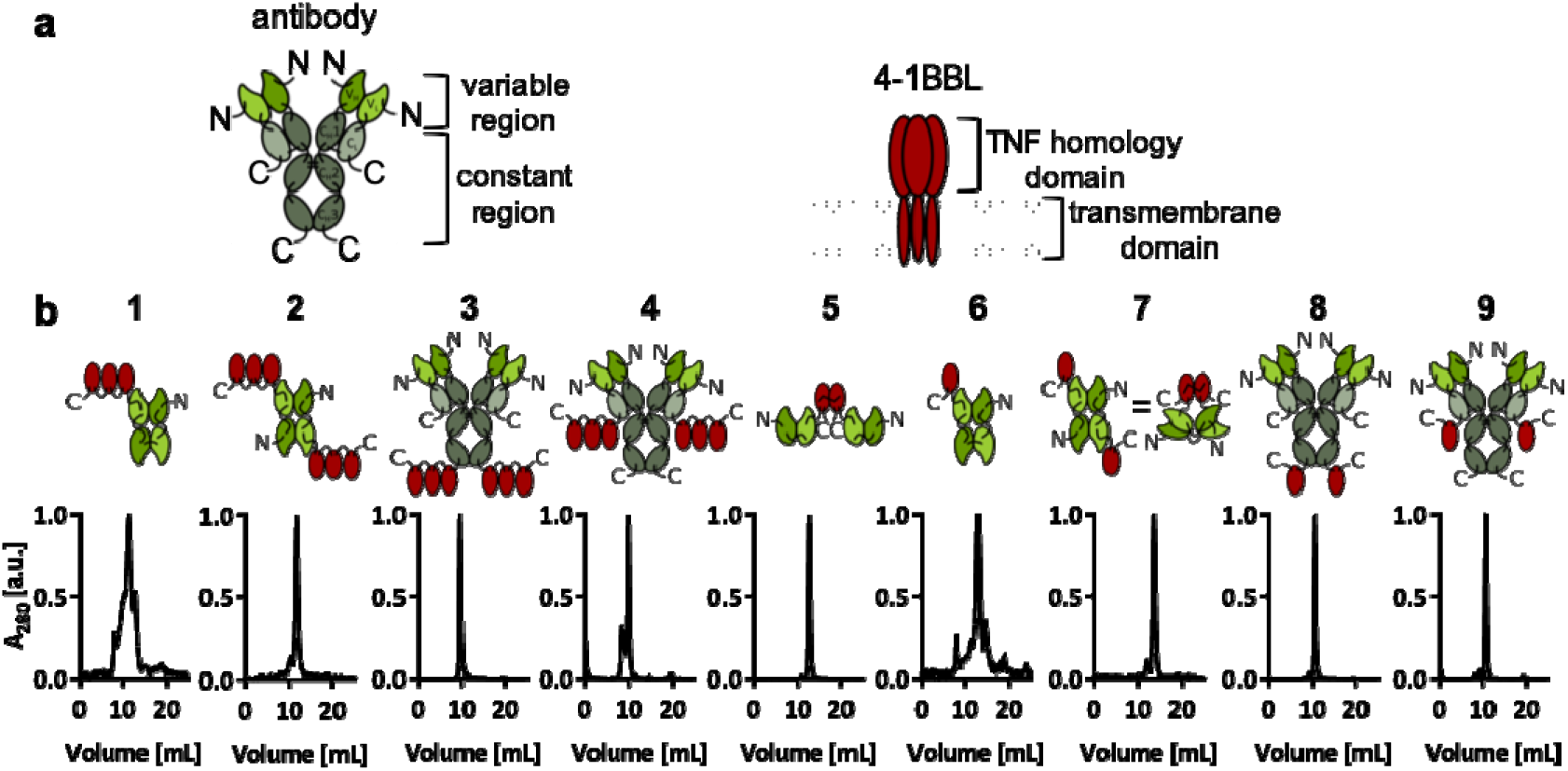
The nine F8-4-1BBL fusion proteins that were designed and tested in this study **(a)** schematic depiction of an antibody in the IgG format and of the human 4-1BBL. The human 4-1BBL is a transmembrane protein which forms a non-covalent homotrimer(57) **(b)** the nine different F8-4-1BBL formats are schematically depicted and size exclusion chromatograms are provided.

**Figure 2** presents a comparative analysis of *in vitro* properties of F8-4-1BBL in various formats. The 4-1BBL and F8 moieties were able to recognize the cognate targets in the **2, 3, 5, 7** and **8** formats. Indeed, all proteins bound with high affinity to murine CTLL-2 cells, which are strongly positive for murine 4-1BB (i.e., the 4-1BBL receptor) [**Figure 2a**] and to recombinant EDA domain of fibronectin [**Figure 2b**]. A functional assay with an NK-κB reporter cell line(66) revealed that all fusion proteins preferentially activated downstream signaling events in the presence of the cognate EDA fibronectin antigen, immobilized on a solid support and thus mimicking the tumor environment [**Figure 2c**]. Formats **5** [consisting of two disulfide-linked 4-1BBL monomeric units fused to scFv(F8)] and **8** [in which monomeric units of 4-1BBL were fused at the C-terminal ends of the heavy chains of IgG(F8)] exhibited the best discrimination between low biological activity in solution and high cytokine activity in the presence of antigen. For this reason, formats **5** and **8** were selected for an *in vivo* characterization of their tumor targeting properties. Format **2** was also included in the comparison, since diabody-based antibody cytokine fusion proteins have previously been used for clinical development programs(2, 64).

**Figure 2:**
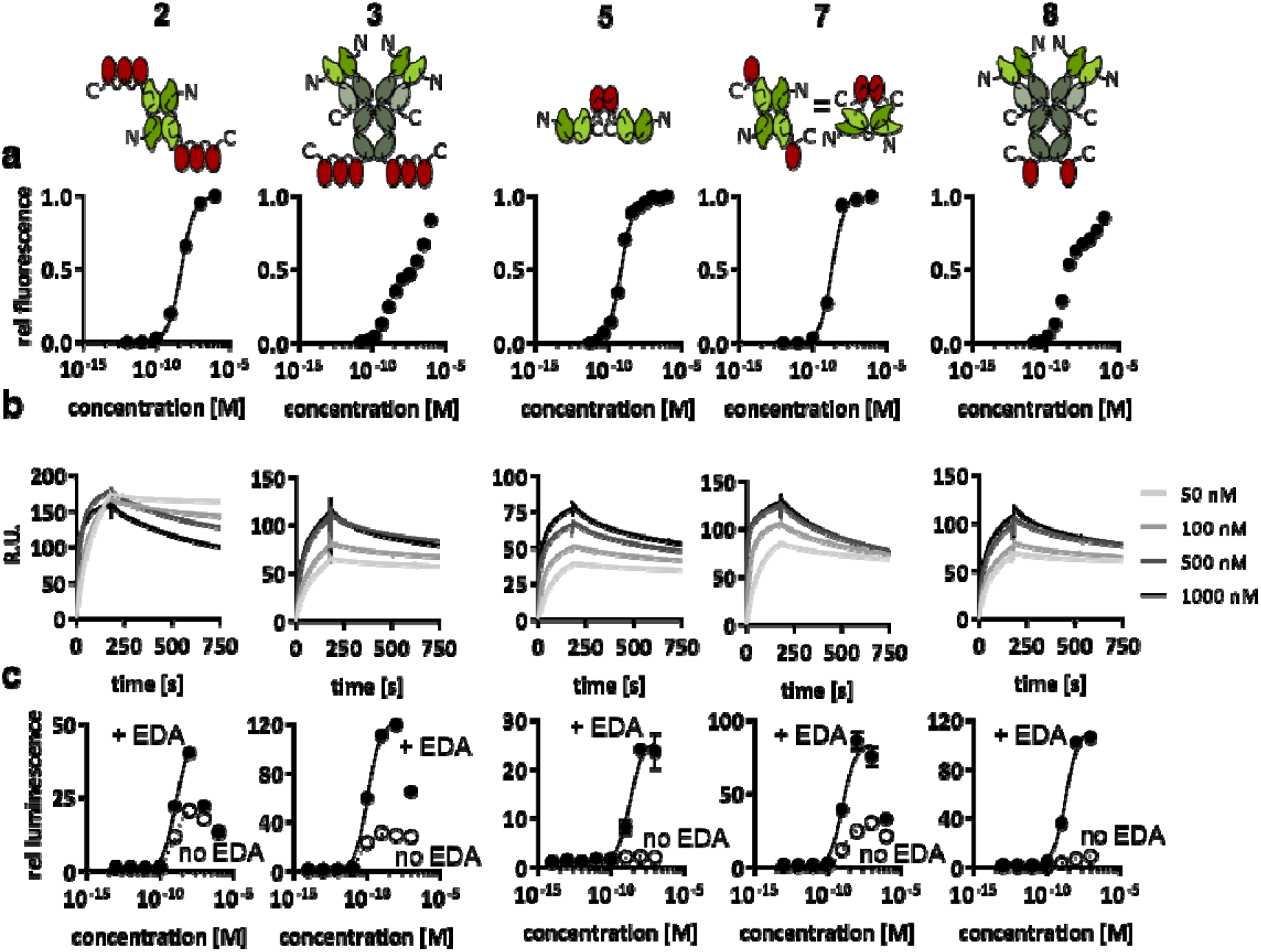
*In vitro* characterization of five F8-4-1BBL formats (a) binding to 4-1BB was measured by flow cytometry with the murine cytotoxic T cell line CTLL-2 which expresses 4-1BB (b) binding of the F8 moiety to the EDA-positive ectodomain of fibronectin was measured by surface plasmon resonance on chips coated with recombinant EDA **(c)** biological activity was tested using an NF-κB reporter cell line that secretes luciferase upon activation of the NF-κB pathway by signaling through 4-1BB. The assay was performed both with and without EDA immobilized on solid support (n = 3). Curve fitting was done using the [agonist] vs response (three parameters) fit of GraphPad Prism v7.0. Data represent mean± SD.

Protein preparations were radioiodinated and injected into immunocompetent 129/Sv mice, bearing subcutaneously-grafted murine F9 teratocarcinomas, which express EDA fibronectin around tumor blood vessels(44). Mice were sacrificed 24 hours after intravenous administration and biodistribution results were expressed as percent of injected dose per gram of tissue [%ID/g] [**Figure 3a**]. Format **2** exhibited only a modest tumor uptake (1.0% ID/g) and poor selectivity. Format 8 showed, as expected, a longer circulatory half-life, as evidenced by the high %ID/g in blood after 24 h, but the tumor uptake and selectivity were not significantly higher compared to KSF-4-1BBL (a fusion protein based on the KSF antibody, specific to hen egg lysozyme and serving as negative control(67)). By contrast, format 5 exhibited a preferential accumulation in the tumor (2.8 % ID/g) and a good tumor-to-normal organ selectivity. EDA targeting was essential for tumor homing, as revealed by the comparison of the biodistribution results with the negative control KSF-4-1BBL fusion protein [**Figure 3a**]. In order to confirm selective tumor uptake with a different methodology, format 5 was injected into tumor-bearing mice. An *ex vivo* immunofluorescence analysis revealed a preferential accumulation of format 5 around tumor blood vessels, while no staining was detectable in normal organs or when the KSF fusion protein was used [**Figure 3b**]. In line with previous reports on this matter(44, 45, 47, 48), the EDA-domain of fibronectin is an ideal target for pharmacodelivery applications in mouse and in man, as the antigen is undetectable in normal adult tissues, but is strongly expressed in the stroma and around the blood vessels in many different tumor types [**Figure 3c**].

**Figure 3:**
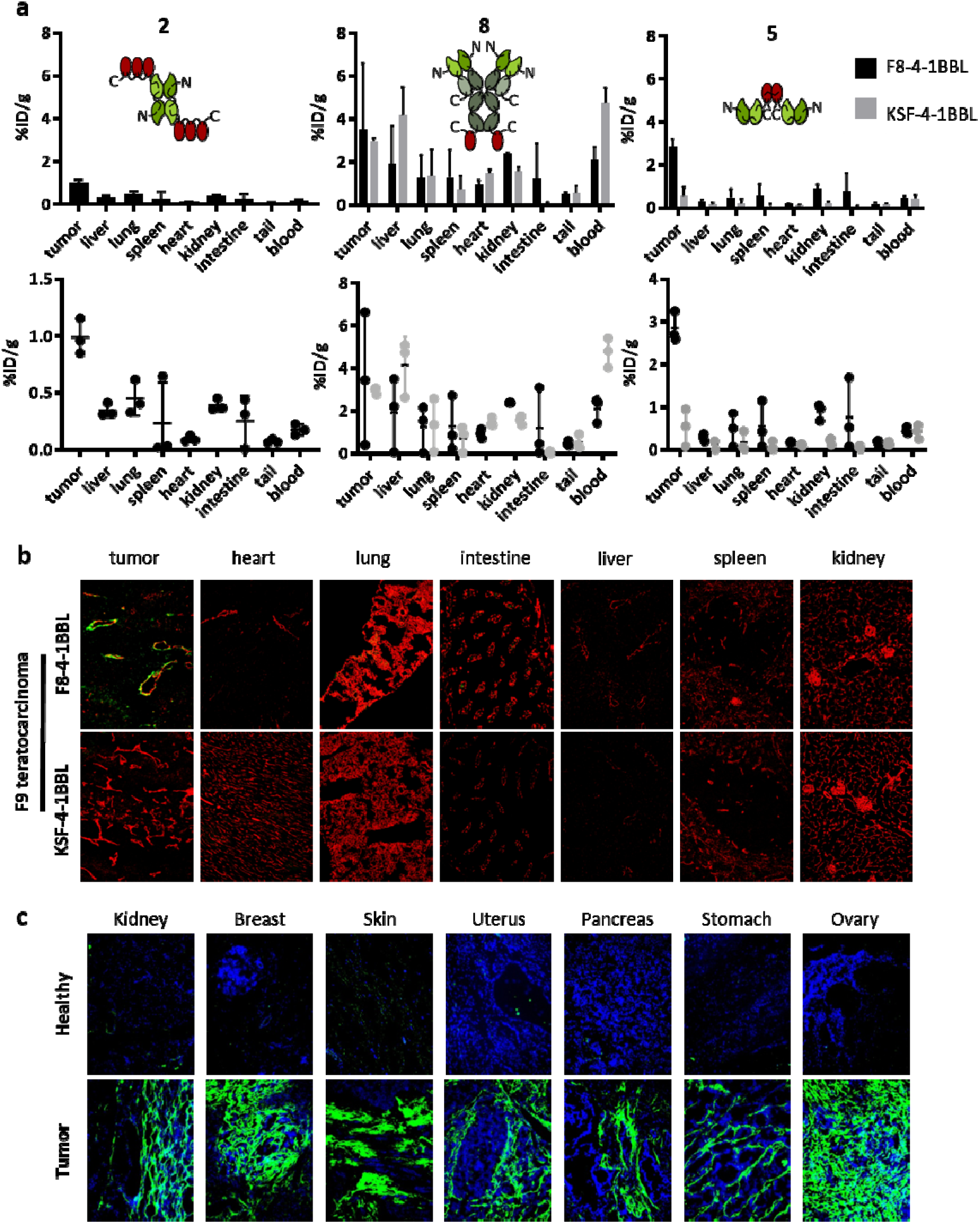
*In vivo* biodistribution studies of three F8-4-1BBL formats (a) The mice were sacrificed 24 h after the injection of the radioiodinated proteins and the radioactivity of excised organs was measured and expressed as percent injected dose per gram of tissue (%ID/g ± SD, n=3). The KSF antibody targeting hen egg lysozyme was used as untargeted control(67). (b) The mice were sacrificed 24 h after the injection of FITC-labelled F8-4-1BBL or KSF-4-1BBL in format 5. The proteins were detected *ex vivo* on cryosections (green: αFITC, red: αCD31). **(c)** The expression of EDA was assessed on human tissue microarrays using FITC-labelled F8 in the IgG format, (green: EDA, blue: nuclei).

Therapy studies were performed using format 5 of F8-4-1-BBL, both in a preventive setting starting at a tumor volume of 40 mm^3^ and in a therapeutic setting starting at a tumor volume of 80 - 100 mm^3^. In a preventive setting in WEHI-164 fibrosarcoma, three out of five mice rejected the tumor using F8-4-1-BBL as single agent, while four out of five mice showed a complete response when treated with PD-1 blockade, alone or in combination with F8-4-1-BBL [**Figure 4a**]. The cured mice rejected subsequent challenges with WEHI-164 fibrosarcoma cells. In some cured mice, a challenge with CT26 colon carcinoma cells was also rejected, similar to what we had previously reported for other F8-based immunocytokine therapeutics (68, 69)[**Supplementary Figure 4**], When the therapy was repeated in mice bearing larger WEHI-164 fibrosarcoma tumors, a significant [p = 0.0427, regular two-way ANOVA, Tukey’s multiple comparison test, day 13] tumor growth retardation was observed in mice treated with F8-4-1BBL [**Figure 4b**]. There was no difference in tumor growth between mice receiving injections of saline and the KSF fusion proteins, underlining the importance of the antigen-dependent activation of 4-1BBL [**Figure 4b**]. Similar experiments performed in immunocompetent mice bearing CT26 tumors showed a tumor regression in 4/5 mice treated with F8-4-1BBL. One mouse was cured, while tumors eventually regrew in the other mice. Therapy was potent also when F8-4-1BBL was combined with PD-1 blockade [**Figure 4c**]. All treatments in all experiments were well tolerated, as indicated by the absence of body weight loss [**Figure 4**].

**Figure 4:**
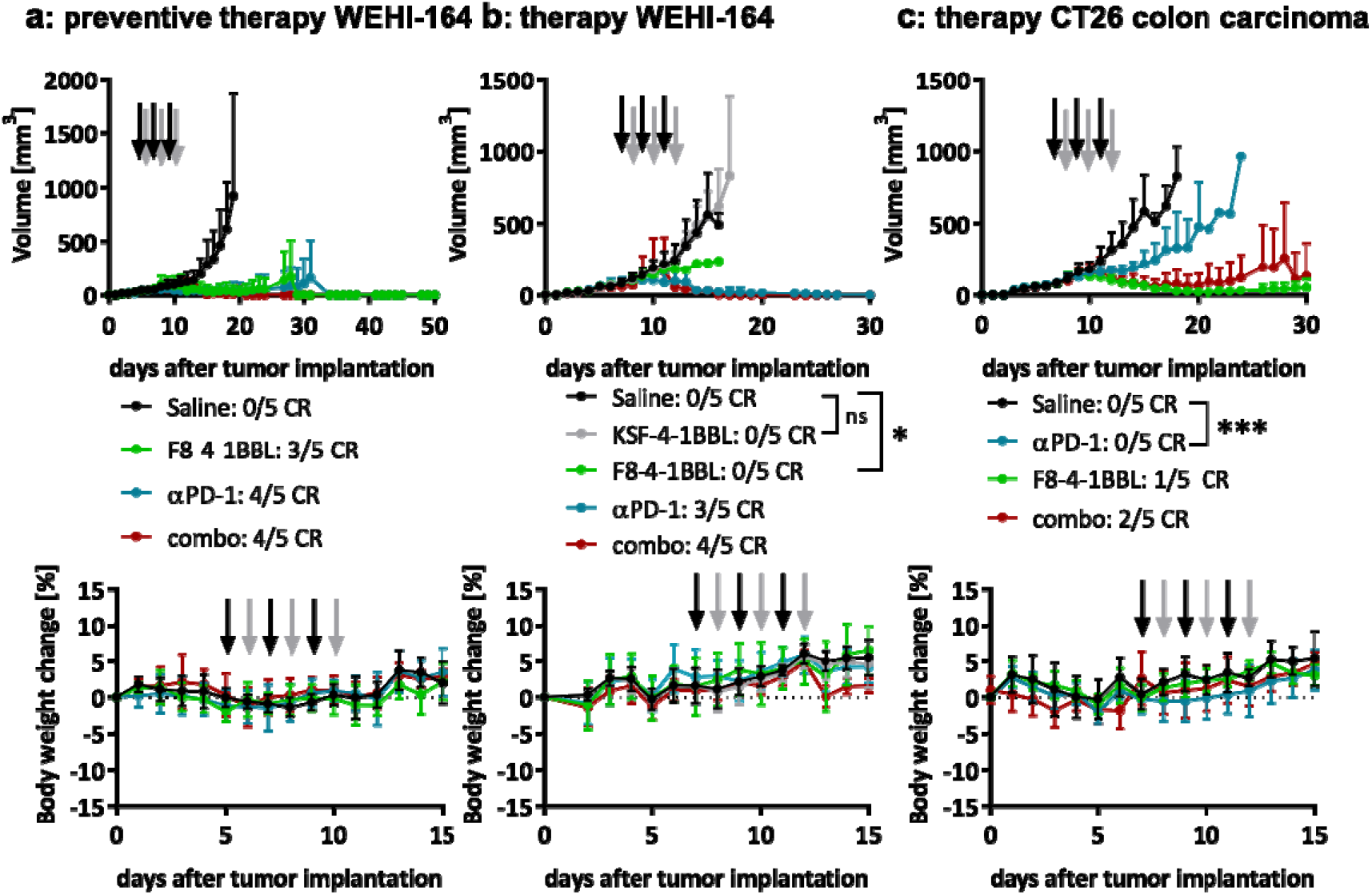
Therapy studies with F8-4-1BBL in format **5 (a)** The preventive therapy in WEHI-164 fibrosarcoma-bearing mice was started on day 5 when the tumors reached a volume of 40 mm^3^. The tumor sizes are shown as mean + SD (n=5). The body weight data is represented as mean body weight change ± SD for each group. (b) The therapy in WEHI-164 fibrosarcoma-bearing mice was started on day 7 when the tumor volume was > 80mm^3^. The tumor sizes are shown as mean + SD (n=5). The statistical results of a regular two-way ANOVA followed by a Tukey’s multiple comparison test using GraphPad Prism v8.4.1 on day 13 are shown (ns: not significant, * p = 0.0427). The body weight data is represented as mean body weight change ± SD for each group. (c) In CT26-colon carcinoma-bearing mice the therapy was started on day 7 when the tumor volume exceeded 80 mm^3^. The tumor sizes are shown as mean + SD (n=5). The result of a regular two-way ANOVA followed by a Tukey’s multiple comparison test using GraphPad Prism v8.4.1 is shown for day 13 (*** p = 0.0004). The body weight data is represented as mean body weight change ± SD for each group, (black arrows: injections of the single-agents, grey arrows: injection of F8-4-1BBL in the combination treatment, CR: complete response)

In order to analyze the tumor infiltrating leukocytes, mice were sacrificed 48 h after the second cycle of injections. Tumors and tumor-draining lymph nodes were excised, homogenized and stained for analysis by flow cytometry. CT26 tumors were found to be highly infiltrated by lymphocytes, in keeping with previous reports(70–72), while WEHI-164 lesions were rather immunologically “cold” [**Figure 5a**]. The proportion of CD8^+^ T cells, specific to AH1 (a retroviral antigen, which plays a dominant role for the rejection of tumors implanted in BALB/c mice (69, 73)) was higher in CT26 tumors [**Figure 5a**]. Treatment with F8-4-1BBL led to a significant increase in intratumoral CD3+ T cell density in both models, but the proportion of CD4+ or CD8+ T cells did not vary substantially. No difference was observed in terms of regulatory T cell (T_reg_) density [**Figure 5a**]. In keeping with what previously reported for other studies(74), the proportion of AH1-specific CD8+ T cells did not vary substantially as a result of pharmacological treatment [**Figure 5a**]. Treatment with F8-4-1BBL led to a decrease in CD3+ and CD4+ T cells in the tumor-draining lymph nodes with a concomitant increase of antigen-presenting cells in CT26 tumor-bearing mice (but not in WEHI-164) [**Figure 5b**]. An increase in the proportion of effector T cells (CD44+CD62L-) was observed among the AH1-specific CD8+ T cells in the tumor-draining lymph nodes [**Figure 5c**]. Virtually all tumor-infiltrating CD8+ T cells were positive for the exhaustion markers PD-1 and CD39 [**Figure 5c**](75, 76). The gating strategy used in the study can be found in **Supplementary Figure 5**. Collectively, the markers used in this study did not detect a phenotypic change in tumor-infiltrating T cells, but an increase in effector T cells was observed for the AH1-specific CD8+ T cell population in tumor-draining lymph nodes as a result of F8-4-1BBL treatment.

**Figure 5:**
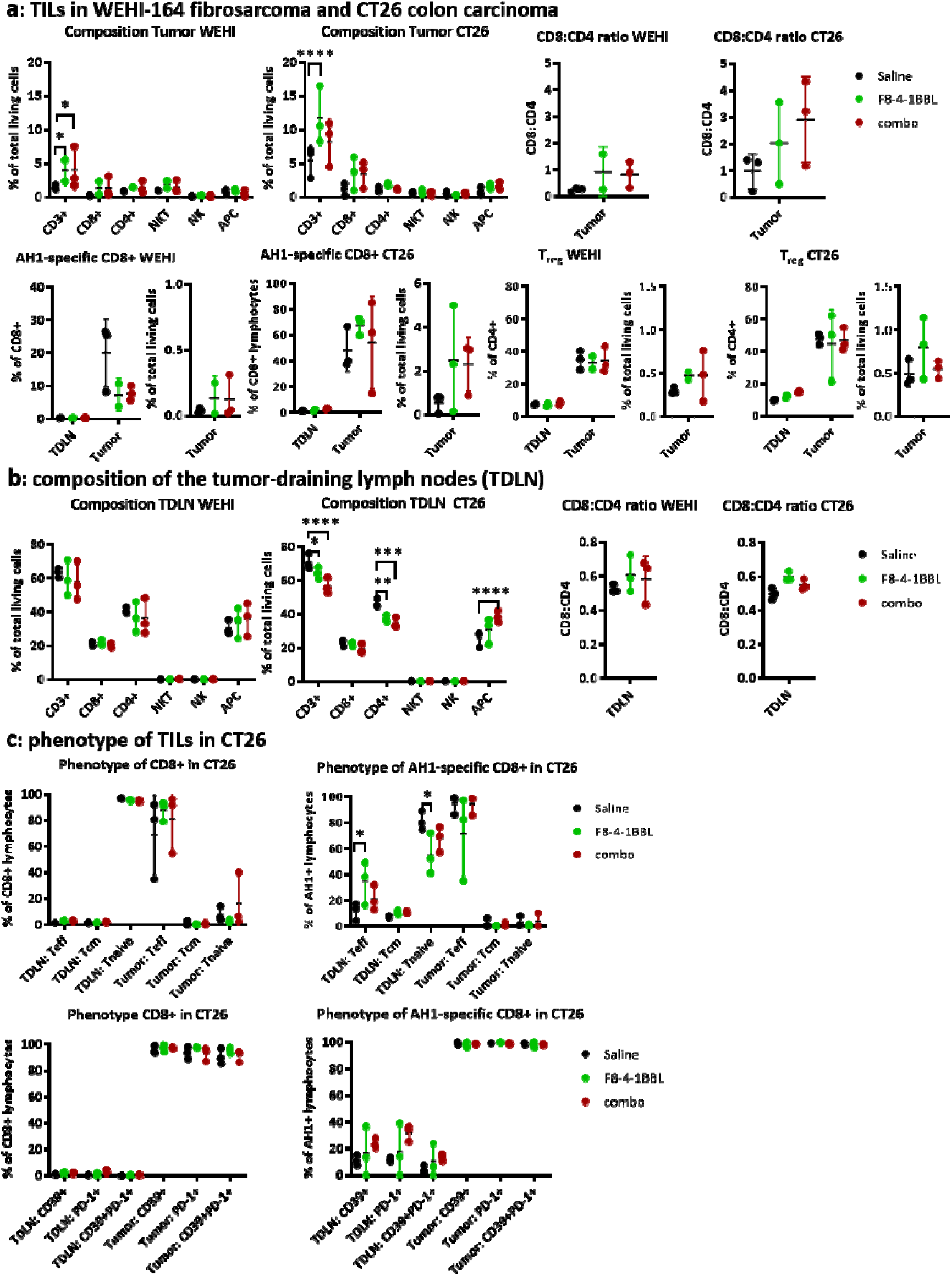
Analysis of tumor-infiltrating leukocytes (TIL) and tumor-draining lymph nodes (TDLN) **(a)** Composition of the tumor-infiltrating immune cells, including the CD8:CD4 ratio, the proportion of AH1-specific CD8+ T cells and regulatory CD4+ T cells in WEHI-164 and CT26 tumors treated with saline, F8-4-1BBL and the combo therapy (αPD-1 and F8-4-1BBL). Proportion of AH1-specific CD8+ T cells and regulatory CD4+ T cell are shown also for matching TDLNs. **(b)** Composition of the TDLNs in WEHI-164 and CT26 tumor-bearing mice, including the CD8:CD4 ratio **(c)** Phenotype of the CD8+ T cells and AH1-specific CD8+ T cells in CD26 tumor-bearing mice from different treatment groups. The phenotype was assessed based on the expression of CD62L, CD44 and the exhaustion markers CD39 and PD-1. The data represents individual values, means and standard deviations. Statistical evaluations were performed using a regular two-way ANOVA followed by a Tukey’s multiple comparison test using GraphPad Prism v8.4.1 [* p < 0.05, ** p < 0.01, *** p < 0.001, **** p < 0.0001] (TIL: tumor-infiltrating leukocyte, NKT: natural killer T cell, NK: natural killer cell, APC: antigen-presenting cell, T_reg_: regulatory T cell, TDLN: tumor-draining lymph node, Teff: effector T cell [CD44+CD62L-], Tcm: central memory T cell [CD44+CD62L+], Tnaive: naïve T cell [CD44-CD62L+])

## DISCUSSION

We have described the development of an antibody-cytokine fusion protein targeted to the tumor neovasculature, featuring an engineered homodimeric 4-1BBL moiety as immunostimulatory payload. The selected format 5 was inactive in solution but regained activity upon clustering on the antigen. Favorable tumor-targeting results and potent tumor growth inhibition were observed *in vivo*, making F8-4-1BBL a promising prototype of for the development of next-generation immunocytokines with “activity-on-demand” properties.

4-1BB, the receptor for 4-1BBL has been recognized as important target for the immunotherapy of cancer, as this member of the TNF receptor superfamily delivers costimulatory signals to activated cytotoxic T cells(77). The first 4-1BB agonistic antibody, urelumab, showed promising anti-cancer activity in preclinical models, but unfortunately revealed substantial hepatotoxicity in clinical trials(78). The hepatic toxicity was mainly due to the activation of liver Kupffer cells and monocytes, leading to a massive infiltration by T cells(78, 79). Efforts are being made to develop 4-1BB agonists with more favorable toxicity profiles that retain potent costimulatory capacities(80–82). In addition to the optimization of anti-4-1BB immunoglobulins(80, 81), various formats of targeted 4-1BB agonists are being investigated. Bispecific antibodies capable of simultaneous recognition of 4-1BB and of tumor-associated antigens (e.g., EGFR or CEA) have been developed and tested in preclinical models of cancer, with encouraging results (82, 83). Novel formats of targeted 4-1BB agonists have recently been considered for clinical development. A FAP-targeted immunocytokine with trimeric single-chain 4-1BBL has recently started phase I clinical testing in cancer patients, but the product did not exhibit a conditional activation of the 4-1BBL moiety(84). A fusion protein of trastuzumab with a 4-1BB-specific anticalin™ has been described(85), which had shown antigen-dependent modulation of 4-1BB agonistic activity *in vitro* and which has recently started clinical trials(85).

The search for antibody-cytokine products with “activity-on-demand” has been recognized as an important research goal, in order to generate products with improved activity and safety profiles(86, 87). One possible strategy features the use of cytokine-binding polypeptides, acting as proteolytically-cleavable inhibitory moieties(88). Fusing cytokines at the C-terminal end of the IgG light chain may restrict conformational changes in the hinge region and slightly modulate cytokine activity upon antibody binding to the cognate antigen(87). The attenuation of cytokine potency by targeted mutagenesis has been considered as a strategy to increase the dose of antibody-cytokine fusion proteins(89) or to conditionally activate tumor cells which express both a tumor-associated antigen and a cytokine receptor (e.g., IFNα receptor) on their surface(90, 91). In addition, the targeted reconstitution of antibodies fused with “split-cytokine” moieties (i.e., subunits of heterodimeric cytokines that can reassemble at the tumor site) has been reported. Until now, the performance of that approach has been limited by the fact that the cytokine subunits used in the study (e.g., the p35 chain of IL12) retained biological activity(92).

The approach described in this article should be generally applicable to members of the TNF superfamily(60). Alternative approaches may involve bispecific antibodies(93, 94), the modular use of small protein domains(85, 95) or of chemically-modified bicyclic peptides(96). Members of the TNF receptor superfamily are particularly suited for cooperative activation strategies in view of their homotrimeric structure and clustering-driven activation properties(97–99). The development of immunotherapeutics with “activity-on-demand” for monomeric cytokines may be more challenging, as one cannot rely on protein assembly for the reconstitution of biological activity.

## MATERIALS AND METHODS

### Cell lines

The murine cytotoxic T cell line CTLL-2 (ATCC^®^ TIB-214), the murine F9 teratocarcinoma cell line (ATCC^®^ CRL-1720), the murine WEHI-164 fibrosarcoma cell line (ATCC^®^ CRL-1751) and the murine CT26 colon carcinoma cell line (ATCC^®^ CRL-2638) were obtained from ATCC, expanded and stored as cryopreserved aliquots in liquid nitrogen. The CTLL-2 cells were grown in RPMI 1640 (Gibco, #21875034) supplemented with 10% FBS (Gibco, #10270106), 1 X antibiotic-antimycoticum (Gibco, #15240062), 2 mM ultraglutamine (Lonza, #BE17-605E/U1), 25 mM HEPES (Gibco, #15630080), 50 μM β-mercaptoethanol (Sigma Aldrich) and 60 U/mL human IL-2 (Proleukin, Roche Diagnostics). The F9 teratocarcinoma cells were grown in DMEM (Gibco, high glucose, pyruvate, #41966-029) supplemented with 10% FBS (Gibco, #10270106) and 1 X antibiotic-antimycoticum (Gibco, #15240062) in flasks coated with 0.1% gelatin (Type B from Bovine Skin, Sigma Aldrich, #G1393). The WEHI-164 fibrosarcoma and the CT26 colon carcinoma were grown in RPMI 1640 (Gibco, #21875034) supplemented with 10% FBS (Gibco, #10270106) and 1 X antibiotic-antimycoticum (Gibco, #15240062). The cells were passaged at the recommended ratios and never kept in culture for more than one month.

### Mice

Eight weeks old female Balb/c and 129/Sv mice were obtained from Janvier. After at least one week of acclimatization, 10^7^ F9 cells, 2.5 x 10^6^ WEHI-164 cells or 4 x 10^6^ CT26 cells were subcutaneously implanted into the right flank. The tumor size was monitored daily by caliper measurements and the volume was calculated using the formula [length x width x width x 0.5]. The animal experiments were carried out under the project license ZH04/2018 granted by the Veterinäramt des Kantons Zürich, Switzerland, in compliance with the Swiss Animal Protection Act (TSchG) and the Swiss Animal Protection Ordinance (TSchV).

### Cloning

A soluble single-chain trimer of murine 4-1BBL was designed by linking the TNF homology domain (amino acids 139 – 309) with a single glycine as a linker. The genetic sequence was ordered from Eurofins Genomics. The sequence was then introduced into a vector encoding the F8 in a diabody format by Gibson Isothermal Assembly. To clone the single-chain variable Fragment (scFv) linked to the 4-1BBL monomer, the genetic sequence encoding the diabody was replaced by the sequence encoding the scFv and two domains of 4-1BBL were removed by PCR followed by blunt-end ligation. Additional base pairs of 4-1BBL were added to the 4-1BBL sequence by PCR followed by blunt-end ligation. The IgG fusion proteins were cloned by fusing the 4-1BBL sequence to the sequence of the antibody in the IgG format by PCR before introducing it into an appropriate vector by restriction cloning. The protein sequences are provided in [**Supplementary Table 1**].

### Protein production

Proteins were produced by transient transfection of CHO-S cells and purified by protein A affinity chromatography as described previously (68, 100, 101). Quality control of the purified products included SDS-PAGE and size exclusion chromatography using an Äkta Pure FPLC system (GE Healthcare) with a Superdex S200 10/300 increase column at a flow rate of 0.75 mL/min (GE Healthcare) [**Figure 1 and Supplementary Figure 1**].

### Binding measurements by Surface Plasmon Resonance

To evaluate the binding kinetics of the F8 antibody fragment to EDA, a CM5 sensor chip (GE Healthcare) was coated with approximately 500 resonance units of an EDA-containing recombinant fragment of fibronectin. The measurements were carried out with a Biacore S200 (GE Healthcare). The contact time was set to 3 min at a flow rate of 20 μL/min followed by a dissociation for 10 min and a regeneration of the chip using 10 mM HCl.

### Binding measurements by Flow Cytometry

In order to measure the binding of the 4-1BBL moiety to cells expressing 4-1BB, CTLL-2 cells were incubated with varying concentrations of the fusion proteins for 1 h. The bound protein was detected by addition of an excess of AlexaFluor488-labelled protein A (Thermofisher, #P11047) and subsequent measurement of the fluorescence using a Cytoflex Flow Cytometer. The mean fluorescence was normalized and the resulting binding curve was fitted using the the [Agonist] vs. response (three parameters) fit of the GraphPad Prism 7.0 a software to estimate the functional K_D_.

### NF-κB response assay

The development of the CTLL-2 reporter cell line is described elsewhere(66). CTLL-2_NF-κB reporter cells were starved by washing the cells twice with prewarmed HBSS (Gibco, #14175095) followed by growth in the absence of IL-2 for 6 - 9 h in RPMI 1640 (Gibco, # 21875034) medium supplemented with 10% FBS (Gibco, #10270106), 1 X antibiotic-antimycoticum (Gibco, # 15240062), 2 mM Ultraglutamine (Lonza, # BE17-605E/U1), 25 mM HEPES (Gibco, # 15630080) and 50 μM β-mercaptoethanol (Sigma Aldrich) in order to reduce the background signal. To coat the wells with antigen, 100 μL 100 nM 11-A-12 fibronectin in phosphate buffered saline (PBS) was added to each well and the plate was incubated at 37°C for 90 min. Cells were seeded in 96-well plates (50,000 cells/well) and growth medium containing varying concentrations of the antibody-cytokine conjugate was added. The cells were incubated at 37°C, 5% CO_2_ for several hours. To assess luciferase production, 20 μL of the supernatant was transferred to an opaque 96-well plate (PerkinElmer, Optiplate-96, white, #6005290) and 80 μL 1 μg/mL Coelenterazine (Carl Roth AG, #4094.3) in phosphate buffered saline (PBS) was added. Luminescence at 595 nm was measured immediately. The relative luminescence was calculated by dividing the obtained results by the results obtained when no inducer was added. The data was fitted using the [Agonist] vs. response (three parameters) fit of the GraphPad Prism 7.0 a software to estimate the EC_50_.

### Quantitative biodistribution studies

Quantitative biodistribution experiments were carried out as described previously(44). Briefly, 8 weeks old female 129/Sv mice were injected subcutaneously in the right flank with 10^7^. F9 teratocarcinoma cells. The tumor size was measured daily with a caliper and the volume was calculated using the formula [volume = length x width x width x 0.5]. When the tumors reached a volume of 100 – 300 mm^3^, 10 μg of radioiodinated protein was injected into the lateral tail vein. The mice were sacrificed 24 h after the injection and the organs were excised and weighed. The radioactivity of the different organs was measured (Packard Cobra II Gamma Counter) and expressed as percentage of injected dose per gram of tissue (%ID/g±SD, *n* = 3).

### *Ex vivo* detection of fluorescently labelled immunocytokines

For fluorescent labelling, the proteins were resuspended in a 0.1 M sodium carbonate buffer at pH 9.1 and an excess of Fluorescein Isothiocyanate (FITC) was added. The reaction was carried out overnight at 4°C. The labelled proteins were separated from unconjugated FITC by PD-10. Approximately 100 μg of fluorescently-labelled protein was injected into the lateral tail-vein of tumor-bearing mice. The mice were sacrificed 24 h after the injection. The organs were excised and embedded in NEG-50 cryoembedding medium (ThermoFisher, Richard-Allan-Scientific, #6502) prior to freezing. For staining, 8 μm cryosections were fixed in acetone and incubated with goat-anti-mouse CD31 (R&D system, #AF3628, 1:200) and rabbit-anti-FITC (Biorad, #4510-7804) followed by donkey-anti-goat-AF594 (Invitrogen, #A11058) and donkey-anti-rabbit-AF488 (Invitrogen, #A21206). Images were acquired using a Zeiss Axioscope 2 mot plus with an Axiocam 503 camera at a 200 X magnification in the RGB mode. The images were processed using the software ImageJ v1.52k setting the thresholds for the red channel to 14-80 and the green channel to 15 −100.

### Immunofluorescence on tissue microarray

Immunofluorescence was performed onto Frozen Tumor and Normal Tissue Array (Biochain, #T6235700). The array was fixed by ice-cold aceton for 5 minutes. After fixation, sections were let dry at room temperature for 10 minutes and then blocked for 45 min with 20% fetal bovine serum in PBS. FITC labeled IgG(F8) was added at 5 μg/ml in 2% BSA/PBS solution for 1h at room temperature. The tissue array was then washed twice with PBS and secondary rabbit anti-FITC antibody (Biorad, #4510-7804) was added to a final 1:1000 dilution in 2% BSA/PBS at room temperature for 1h. After washing the array twice with PBS, Goat Anti-Rabbit Alexa-488 (ThermoFisher, #A11032) was added to a final 1:500 dilution in 2% BSA/PBS. Dapi was used to counterstain nuclei. Slides were analyzed with Axioskop2 plus microscope (Zeiss).

### Therapy studies

After the subcutaneous implantation of the tumor cells into the right flank of 8 weeks old female mice, the tumor size was monitored by caliper measurements on a daily basis [volume = length x width x width x 0.5]. In the preventive setting, the therapy was started when the tumor reached a volume of 40 mm^3^ and for the therapeutic setting, the therapy was started when the tumors reached a volume of 80 – 100 mm^3^. The mice were grouped in order to obtain groups of similar average tumor size (n = 5). The mice either received 100 μL Saline (PBS, Gibco, #1010023), 500 μg F8(scFv)-4-1BBL, 200 μg αPD-1 (BioXCell, clone 29F.1A12) or a combination of the checkpoint inhibitor and the immunocytokine. For the combination treatment, the checkpoint inhibitor was administered one day prior to the immunocytokine. The therapeutic agents were administrated every second day in a total of three cycles intravenously into the lateral tail vein. The animals were sacrificed if the tumor diameter exceeded 15 mm or when the tumor started to ulcerate. Some cured mice were rechallenged by the subcutaneous injection of WEHI 164 or CT26 tumor cells after being tumor-free for at least 4 weeks. Statistical evaluations were done using a standard two-way ANOVA followed by the Tukey’s multiple comparison test with GraphPad Prism v8.4.1.

### Analysis of tumor-infiltrating lymphocytes by flow cytometry

The mice were sacrificed 48 h after the second therapy cycle. The tumor-draining lymph nodes as well as the tumor were excised. A single-cell suspension of the tumor was obtained by digesting it in RPMI 1640 supplemented with 1 mg/mL collagenase II and 100 μg/mL DNase I for 30 min at 37°C. After the digestion, the suspension was passed through a 70 μm cell strainer. If necessary, the red blood cells were removed using a red blood cell lysis buffer (Roche). The lymph nodes were smashed on a 70 μm cell strainer and washed with PBS. For cell surface staining, cells were incubated with a mix of suitable antibodies: αCD3-APC/Cy7 (Biolegend, #100222), αCD4-APC (Biolegend, #100412), αCD8-FITC (Biolegend, #100706), αNK1.1-PE (Biolegend, #108708), αCD62L-BV421 (Biolegend, #104436), αCD44-APC/Cy7 (Biolegend, #103028), αMHCII(IA/IE)-BV421 (Biolegend, #107631), αPD-1-BV421 (Biolegend, #109121) and αCD39-APC (Biolegend, #143809). After staining of the cell surface markers, the cells were stained with 7-AAD (Biolegend) for live/dead discrimination. For intracellular staining, the cells were first stained with Zombie Red (SigmaAldrich) and then the cell surface stain was performed. The cells were fixed and permeabilized using the eBioscience^™^ FoxP3/Transcription Factor Staining Buffer Set (Thermofisher, #00-5523-00) according to the manufacturer’s instructions. The fluorescence was measured using a Cytoflex Flow Cytometer and the data was evaluated using the FlowJo software. The gating strategy is depicted in Supplementary Figure 5. Statistical evaluations were done using a regular two-way ANOVA followed by a Tukey’s multiple comparison test or a regular oneway ANOVA followed by a Sidak’s multiple comparison test in GraphPad Prism v8.4.1.

## ACKNOWLEDGEMENTS

Financial support by the ETH Zürich, the Swiss National Science Foundation (grant number 310030_182003/1), the European Research Council (ERC) under the European Union’s Horizon 2020 research and innovation program (grant agreement 670603), and the Federal Commission for Technology and Innovation (KTI, grant number 12803.1 VOUCH-LS) is gratefully acknowledged. In addition, Sabrina Müller and Fiona Amman are gratefully acknowledged for their technical assistance especially for the protein production. The authors would also like to thank Lisa Nadal for her help with some immunofluorescence stainings.

## AUTHOR CONTRIBUTIONS

D.N. and J.M. designed and planned the study. J.M. performed most of the experiments. M.S. helped with the biodistribution studies with radiolabeled proteins. M.S. designed the infiltrate analysis and helped to perform the experiment. A.V. coordinated immunohistochemical studies. J.M. prepared the figures. D.N. and J.M. wrote the manuscript.

## DATA AVAILABILITY STATEMENT

The datasets generated during and/or analyzed during the current study are available from the corresponding author on reasonable request.

