## SupplementaryData for "An engineered 4-1BBL fusion protein with “activity-on-demand”" (from another version of this preprint: version 1, 2020-04-30)

**F8(scDb)-(4-1BBL)$_3$**: F8_VH-*linker*-F8_VL-*linker*- F8_VH-*linker*-F8_VL-*linker*-4-1BBL-*linker*-4-1BBL-*linker*-4-1BBL (Format **1**)
EVQLLESGGGLVQPGGSLRLSCAASGFTFSLFTMSWVRQAPGKGLEWVSAISGSGGSTYYADSVKGRFTISRDNSKNTLYLQ
MNSLRAEDTAVYYCAKSTHLYLFDYWGQGTLVTVSS-*GGSGG*-EIVLTQSPGTLSLSPGERATLSCRASQSVSMPFLAWYQQ
KPGQAPRLLIYGASSRATGIPDRFSGSGSGTDFTLTISRLEPEDFAVYYCQQMRGRPPTFGQGTKVEIK-*GGGGSGGGGSGG*
*GGS*-EVQLLESGGGLVQPGGSLRLSCAASGFTFSLFTMSWVRQAPGKGLEWVSAISGSGGSTYYADSVKGRFTISRDNSKNT
LYLQMNSLRAEDTAVYYCAKSTHLYLFDYWGQGTLVTVSS-*GGSGG*-EIVLTQS PGTLSLSPGERATLSCRASQSVSMPFLA
WYQQKPGQAPRLLIYGASSRATGIPDRFSGSGSGTDFTLTISRLEPEDFAVYYCQQMRGRPPTFGQGTKVEIK-*SSSSGSSSS*
*GSSSSG*-ATTQQGSPVFAKLLAKN QASLCNTTLNWHSQDGAGSSYLSQGLRYEEDKKELVVDSPGLYYVFLELKLSPTFTNT
GHKVQGWVSLVLQAKPQVDDFDNLALTVELFPCSMENKLVDRSWSQLLLLKAGHRLSVGLRAYLHGAQDAYRDWELSYP
NTTSFGLFLVKPDNPWE-*G*-ATTQQGSPVFAKLLAKNQASLCNTTLNWHSQDGAGSSYLSQGLRYEEDKKELVVDSPGLYY
VFLELKLSPTFTNTGHKVQGWVSLVLQAKPQVDDFDNLALTVELFPCSMENKLVDRSWSQLLLLKAGHRLSVGLRAYLHGA
QDAYRDWELSYPNTTSFGLFLVKPDNPWE-*G*-ATTQQGSPVFAKLLAKNQASLCNTTLNWHSQDGAGSSYLSQGLRYEED
KKELVVDSPGLYYVFLELKLSPTFTNTGHKVQGWVSLVLQAKPQVDDFDNLALTVELFPCSMENKLVDRSWSQLLLLKAGH
RLSVGLRAYLHGAQDAYRDWELSYPNTTSFGLFLVKPDNPWE

**F8(dDb)-(4-1BBL)$_3$**: F8_VH-*linker*-F8_VL-*linker*-4-1BBL-*linker*-4-1BBL-*linker*-4-1BBL (Format **2**)
EVQLLESGGGLVQPGGSLRLSCAASGFTFSLFTMSWVRQAPGKGLEWVSAISGSGGSTYYADSVKGRFTISRDNSKNTLYLQ
MNSLRAEDTAVYYCAKSTHLYLFDYWGQGTLVTVSS-*GGSGG*-EIVLTQSPGTLSLSPGERATLSCRASQSVSMPFLAWYQQ
KPGQAPRLLIYGASSRATGIPDRFSGSGSGTDFTLTISRLEPEDFAVYYCQQMRGRPPTFGQGTKVEIK-*SSSSGSSSSGSSSSG*
-ATTQQGSPVFAKLLAKNQASLCNTTLNWHSQDGAGSSYLSQGLRYEEDKKELVVDSPGLYYVFLELKLSPTFTNTGHKVQG
WVSLVLQAKPQVDDFDNLALTVELFPCSMENKLVDRSWSQLLLLKAGHRLSVGLRAYLHGAQDAYRDWELSYPNTTSFGL
FLVKPDNPWE-*G*-ATTQQGSPVFAKLLAKNQASLCNTTLNWHSQDGAGSSYLSQGLRYEEDKKELVVDSPGLYYVFLELKLS
PTFTNTGHKVQGWVSLVLQAKPQVDDFDNLALTVELFPCSMENKLVDRSWSQLLLLKAGHRLSVGLRAYLHGAQDAYRD
WELSYPNTTSFGLFLVKPDNPWE-*G*-ATTQQGSPVFAKLLAKNQASLCNTTLNWHSQDGAGSSYLSQGLRYEEDKKELVVD
SPGLYYVFLELKLSPTFTNTGHKVQGWVSLVLQAKPQVDDFDNLALTVELFPCSMENKLVDRSWSQLLLLKAGHRLSVGLRA
YLHGAQDAYRDWELSYPNTTSFGLFLVKPDNPWE

**F8(IgG)-(4-1BBL)$_3$_HC**: VL-CL** VH-CH-*linker*-4-1BBL-*linker*-4-1BBL-*linker*-4-1BBL (Format **3**)
EIVLTQSPGTLSLSPGERATLSCRASQSVSMPFLAWYQQKPGQAPRLLIYGASSRATGIPDRFSGSGSGTDFTLTISRLEPEDF
AVYYCQQMRGRPPTFGQGTKVEIKRTVAAPSVFIFPPSDEQLKSGTASVVCLLNNFYPREAKVQWKVDNALQSGNSQESV
TEQDSKDSTYSLSSTLTLSKADYEKHKVYACEVTHQGLSSPVTKSFNRGEC**
EVQLLESGGGLVQPGGSLRLSCAASGFTFSLFTMSWVRQAPGKGLEWVSAISGSGGSTYYADSVKGRFTISRDNSKNTLYLQ
MNSLRAEDTAVYYCAKSTHLYLFDYWGQGTLVTVSSASTKGPSVFPLAPSSKSTSGGTAALGCLVKDYFPEPVTVSWNSGAL
TSGVHTFPAVLQSSGLYSLSSVVTVPSSSLGTQTYICNVNHKPSNTKVDKKVEPKSCDKTHTCPPCPAPELLGGPSVFLFPPKP
KDTLMISRTPEVTCVVVDVSHEDPEVKFNWYVDGVEVHNAKTKPREEQYNSTYRVVSVLTVLHQDWLNGKEYKCKVSNKA
LPAPIEKTISKAKGQPREPQVYTLPPSRDELTKNQVSLTCLVKGFYPSDIAVEWESNGQPENNYKTTPPVLDSDGSFFLYSKLT
VDKSRWQQGNVFSCSVMHEALHNHYTQKSLSLSPGK-*SSSSGSSSSGSSSSG*-ATTQQGSPVFAKLLAKNQASLCNTTLNW
HSQDGAGSSYLSQGLRYEEDKKELVVDSPGLYYVFLELKLSPTFTNTGHKVQGWVSLVLQAKPQVDDFDNLALTVELFPCS
MENKLVDRSWSQLLLLKAGHRLSVGLRAYLHGAQDAYRDWELSYPNTTSFGLFLVKPDNPWE-*G*-ATTQQGSPVFAKLLA
KNQASLCNTTLNWHSQDGAGSSYLSQGLRYEEDKKELVVDSPGLYYVFLELKLSPTFTNTGHKVQGWVSLVLQAKPQVDDF
DNLALTVELFPCSMENKLVDRSWSQLLLLKAGHRLSVGLRAYLHGAQDAYRDWELSYPNTTSFGLFLVKPDNPWE-*G*-ATT
QQGSPVFAKLLAKNQASLCNTTLNWHSQDGAGSSYLSQGLRYEEDKKELVVDSPGLYYVFLELKLSPTFTNTGHKVQGWVS
LVLQAKPQVDDFDNLALTVELFPCSMENKLVDRSWSQLLLLKAGHRLSVGLRAYLHGAQDAYRDWELSYPNTTSFGLFLVK
PDNPWE

**F8(IgG)-(4-1BBL)$_3$_LC**: VL-CL-*linker*-4-1BBL-*linker*-4-1BBL-*linker*-4-1BBL** VH-CH (Format **4**)
EIVLTQSPGTLSLSPGERATLSCRASQSVSMPFLAWYQQKPGQAPRLLIYGASSRATGIPDRFSGSGSGTDFTLTISRLEPEDF
AVYYCQQMRGRPPTFGQGTKVEIKRTVAAPSVFIFPPSDEQLKSGTASVVCLLNNFYPREAKVQWKVDNALQSGNSQESV
TEQDSKDSTYSLSSTLTLSKADYEKHKVYACEVTHQGLSSPVTKSFNRGEC-*SSSSGSSSSGSSSSG*-ATTQQGSPVFAKLLAKN
QASLCNTTLNWHSQDGAGSSYLSQGLRYEEDKKELVVDSPGLYYVFLELKLSPTFTNTGHKVQGWVSLVLQAKPQVDDFD
NLALTVELFPCSMENKLVDRSWSQLLLLKAGHRLSVGLRAYLHGAQDAYRDWELSYPNTTSFGLFLVKPDNPWE-*G*-ATTQ
QGSPVFAKLLAKNQASLCNTTLNWHSQDGAGSSYLSQGLRYEEDKKELVVDSPGLYYVFLELKLSPTFTNTGHKVQGWVSL
VLQAKPQVDDFDNLALTVELFPCSMENKLVDRSWSQLLLLKAGHRLSVGLRAYLHGAQDAYRDWELSYPNTTSFGLFLVKP
DNPWE-*G*-ATTQQGSPVFAKLLAKNQASLCNTTLNWHSQDGAGSSYLSQGLRYEEDKKELVVDSPGLYYVFLELKLSPTFTN
TGHKVQGWVSLVLQAKPQVDDFDNLALTVELFPCSMENKLVDRSWSQLLLLKAGHRLSVGLRAYLHGAQDAYRDWELSY
PNTTSFGLFLVKPDNPWE**
EVQLLESGGGLVQPGGSLRLSCAASGFTFSLFTMSWVRQAPGKGLEWVSAISGSGGSTYYADSVKGRFTISRDNSKNTLYLQ
MNSLRAEDTAVYYCAKSTHLYLFDYWGQGTLVTVSSASTKGPSVFPLAPSSKSTSGGTAALGCLVKDYFPEPVTVSWNSGAL

TSGVHTFPAVLQSSGLYSLSSVVTVPSSSLGTQTYICNVNHKPSNTKVDKKVEPKSCDKTHTCPPCPAPELLGGPSVFLFPPKP
KDTLMISRTPEVTCVVVDVSHEDPEVKFNWYVDGVEVHNAKTKPREEQYNSTYRVVSVLTVLHQDWLNGKEYKCKVSNKA
LPAPIEKTISKAKGQPREPQVYTLPPSRDELTKNQVSLTCLVKGFYPSDIAVEWESNGQPENNYKTTPPVLDSDGSFFLYSKLT
VDKSRWQQGNVFSCSVMHEALHNHYTQKSLSLSPGK

**F8(scFv)-4-1BBL**: F8_VH-*linker*-F8_VL-*linker*-4-1BBL (Format **5**)
EVQLLESGGGLVQPGGSLRLSCAASGFTFSLFTMSWVRQAPGKGLEWVSAISGSGGSTYYADSVKGRFTISRDNSKNTLYLQ
MNSLRAEDTAVYYCAKSTHLYLFDYWGQGTLVTVSS-*GGGGSGGGGSGGGG*-EIVLTQSPGTLSLSPGERATLSCRASQSVS
MPFLAWYQQKPGQAPRLLIYGASSRATGIPDRFSGSGSGTDFTLTISRLEPEDFAVYYCQQMRGRPPTFGQGTKVEIK-*SSSS*
*GSSSSGSSSS*-GATTQQGSPVFAKLLAKNQASLCNTTLNWHSQDGAGSSYLSQGLRYEEDKKELVVDSPGLYYVFLELKLSPT
FTNTGHKVQGWVSLVLQAKPQVDDFDNLALTVELFPCSMENKLVDRSWSQLLLLKAGHRLSVGLRALHGAQDAYRDWEL
SYPNTTSFGLFLVKPDNPWE

**F8(scDb)-4-1BBL**: F8_VH-*linker*-F8_VL-*linker*- F8_VH-*linker*-F8_VL-*linker*-4-1BBL (Format **6**)
EVQLLESGGGLVQPGGSLRLSCAASGFTFSLFTMSWVRQAPGKGLEWVSAISGSGGSTYYADSVKGRFTISRDNSKNTLYLQ
MNSLRAEDTAVYYCAKSTHLYLFDYWGQGTLVTVSS-*GGSGG*-EIVLTQSPGTLSLSPGERATLSCRASQSVSMPFLAWYQQ
KPGQAPRLLIYGASSRATGIPDRFSGSGSGTDFTLTISRLEPEDFAVYYCQQMRGRPPTFGQGTKVEIK-*GGGGSGGGGSGG*
*GGS*-EVQLLESGGGLVQPGGSLRLSCAASGFTFSLFTMSWVRQAPGKGLEWVSAISGSGGSTYYADSVKGRFTISRDNSKNT
LYLQMNSLRAEDTAVYYCAKSTHLYLFDYWGQGTLVTVSS-*GGSGG*-EIVLTQSPGTLSLSPGERATLSCRASQSVSMPFLA
WYQQKPGQAPRLLIYGASSRATGIPDRFSGSGSGTDFTLTISRLEPEDFAVYYCQMRGRPPTFGQGTKVEIK-*SSSSGSSSSGS*
*SSSG*-ATTQQGSPVFAKLLAKNQASLCNTTLNWHSQDGAGSSYLSQGLRYEEDKKELVVDSPGLYYVFLELKLSPTFTNTGHK
VQGWVSLVLQAKPQVDDFDNLALTVELFPCSMENKLVDRSWSQLLLLKAGHRLSVGLRAYLHGAQDAYRDWELSYPNTTS
FGLFLVKPDNPWE

**F8(dDb)-4-1BBL**: F8_VH-*linker*-F8_VL-*linker*-4-1BBL (Format **7**)
EVQLLESGGGLVQPGGSLRLSCAASGFTFSLFTMSWVRQAPGKGLEWVSAISGSGGSTYYADSVKGRFTISRDNSKNTLYLQ
MNSLRAEDTAVYYCAKSTHLYLFDYWGQGTLVTVSS-*GGSGG*-EIVLTQSPGTLSLSPGERATLSCRASQSVSMPFLAWYQQ
KPGQAPRLLIYGASSRATGIPDRFSGSGSGTDFTLTISRLEPEDFAVYYCQQMRGRPPTFGQGTKVEIK-*SSSSGSSSSGSSSSG*
-ATTQQGSPVFAKLLAKNQASLCNTTLNWHSQDGAGSSYLSQGLRYEEDKKELVVDSPGLYYVFLELKLSPTFTNTGHKVQG
WVSLVLQAKPQVDDFDNLALTVELFPCSMENKLVDRSWSQLLLLKAGHRLSVGLRAYLHGAQDAYRDWELSYPNTTSFGL
FLVKPDNPWE

**F8(IgG)-4-1BBL_HC**: VL-CL** VH-CH-*linker*-4-1BBL (Format **8**)
EIVLTQSPGTLSLSPGERATLSCRASQSVSMPFLAWYQQKPGQAPRLLIYGASSRATGIPDRFSGSGSGTDFTLTISRLEPEDF
AVYYCQQMRGRPPTFGQGTKVEIKRTVAAPSVFIFPPSDEQLKSGTASVVCLLNNFYPREAKVQWKVDNALQSGNSQESV
TEQDSKDSTYSLSSTLTLSKADYEKHKVYACEVTHQGLSSPVTKSFNRGEC**
EVQLLESGGGLVQPGGSLRLSCAASGFTFSLFTMSWVRQAPGKGLEWVSAISGSGGSTYYADSVKGRFTISRDNSKNTLYLQ
MNSLRAEDTAVYYCAKSTHLYLFDYWGQGTLVTVSSASTKGPSVFPLAPSSKSTSGGTAALGCLVKDYFPEPVTVSWNSGAL
TSGVHTFPAVLQSSGLYSLSSVVTVPSSSLGTQTYICNVNHKPSNTKVDKKVEPKSCDKTHTCPPCPAPELLGGPSVFLFPPKP
KDTLMISRTPEVTCVVVDVSHEDPEVKFNWYVDGVEVHNAKTKPREEQYNSTYRVVSVLTVLHQDWLNGKEYKCKVSNKA
LPAPIEKTISKAKGQPREPQVYTLPPSRDELTKNQVSLTCLVKGFYPSDIAVEWESNGQPENNYKTTPPVLDSDGSFFLYSKLT
VDKSRWQQGNVFSCSVMHEALHNHYTQKSLSLSPGK-*SSSSGSSSSGSSSSG*-ATTQQGSPVFAKLLAKNQASLCNTTLNW
HSQDGAGSSYLSQGLRYEEDKKELVVDSPGLYYVFLELKLSPTFTNTGHKVQGWVSLVLQAKPQVDDFDNLALTVELFPCS
MENKLVDRSWSQLLLLKAGHRLSVGLRAYLHGAQDAYRDWELSYPNTTSFGLFLVKPDNPWE

**F8(IgG)-4-1BBL_LC**: VL-CL-*linker*-4-1BBL** VH-CH (Format **9**)
EIVLTQSPGTLSLSPGERATLSCRASQSVSMPFLAWYQQKPGQAPRLLIYGASSRATGIPDRFSGSGSGTDFTLTISRLEPEDF
AVYYCQQMRGRPPTFGQGTKVEIKRTVAAPSVFIFPPSDEQLKSGTASVVCLLNNFYPREAKVQWKVDNALQSGNSQESV
TEQDSKDSTYSLSSTLTLSKADYEKHKVYACEVTHQGLSSPVTKSFNRGEC-*SSSSGSSSSGSSSSG*-ATTQQGSPVFAKLLAKN
QASLCNTTLNWHSQDGAGSSYLSQGLRYEEDKKELVVDSPGLYYVFLELKLSPTFTNTGHKVQGWVSLVLQAKPQVDDFD
NLALTVELFPCSMENKLVDRSWSQLLLLKAGHRLSVGLRAYLHGAQDAYRDWELSYPNTTSFGLFLVKPDNPWE **
EVQLLESGGGLVQPGGSLRLSCAASGFTFSLFTMSWVRQAPGKGLEWVSAISGSGGSTYYADSVKGRFTISRDNSKNTLYLQ
MNSLRAEDTAVYYCAKSTHLYLFDYWGQGTLVTVSSASTKGPSVFPLAPSSKSTSGGTAALGCLVKDYFPEPVTVSWNSGAL
TSGVHTFPAVLQSSGLYSLSSVVTVPSSSLGTQTYICNVNHKPSNTKVDKKVEPKSCDKTHTCPPCPAPELLGGPSVFLFPPKP
KDTLMISRTPEVTCVVVDVSHEDPEVKFNWYVDGVEVHNAKTKPREEQYNSTYRVVSVLTVLHQDWLNGKEYKCKVSNKA
LPAPIEKTISKAKGQPREPQVYTLPPSRDELTKNQVSLTCLVKGFYPSDIAVEWESNGQPENNYKTTPPVLDSDGSFFLYSKLT
VDKSRWQQGNVFSCSVMHEALHNHYTQKSLSLSPGK

**KSF(IgG)-4-1BBL_HC**: KSF_VL-KSF_CL** KSF_VH-KSF_CH-*linker*-4-1BBL
SELTQDPAVSVALGQTVRITCQGDSLRSYYASWYQQKPGQAPVLVIYGKNNRPSGIPDRFSGSSSGNTASLTITGAQAEDEA
DYYCNSSPLNRLAVVFGGGTKLTVLGCNSSPLNRLAVVFGGGTKLTVLGQPKAAPSVTLFPPSSEELQANKATLVCLISDFYP
GAVTVAWKADSSPVKAGVETTTPSKQSNNKYAASSYLSLTPEQWKSHKSYSCQVTHEGSTVEKTVAPTECS**

EVQLLESGGGLVQPGGSLRLSCAASGFTFSSYAMSWVRQAPGKGLEWVSAISGSGGSTYYADSVKGRFTISRDNSKNTLYL
QMNSLRAEDTAVYYCAKSPKVSLFDYWGQGTLVTVSSASTKGPSVFPLAPSSKSTSGGTAALGCLVKDYFPEPVTVSWNSG
ALTSGVHTFPAVLQSSGLYSLSSVVTVPSSSLGTQTYICNVNHKPSNTKVDKKVEPKSCDKTHTCPPCPAPELLGGPSVFLFPP
KPKDTLMISRTPEVTCVVVDVSHEDPEVKFNWYVDGVEVHNAKTKPREEQYNSTYRVVSVLTVLHQDWLNGKEYKCKVSN
KALPAPIEKTISKAKGQPREPQVYTLPPSRDELTKNQVSLTCLVKGFYPSDIAVEWESNGQPENNYKTTPPVLDSDGSFFLYSK
LTVDKSRWQQGNVFSCSVMHEALHNHYTQKSLSLSPGK-*SSSSGSSSSGSSSS*-GATTQQGSPVFAKLLAKNQASLCNTTLN
WHSQDGAGSSYLSQGLRYEEDKKELVVDSPGLYYVFLELKLSPTFTNTGHKVQGWVSLVLQAKPQVDDFDNLALTVELFPC
SMENKLVDRSWSQLLLLKAGHRLSVGLRALHGAQDAYRDWELSYPNTTSFGLFLVKPDNPWE

**KSF(scFv)-4-1BBL**: KSF_VH-*linker*-KSF_VL-*linker*-4-1BBL

EVQLLESGGGLVQPGGSLRLSCAASGFTFSSYAMSWVRQAPGKGLEWVSAISGSGGSTYYADSVKGRFTISRDNSKNTLYL
QMNSLRAEDTAVYYCAKSPKVSLFDYWGQGTLVTVSS-*GGGGSGGGGSGGGGS*-SELTQDPAVSVALGQTVRITCQGDSL
RSYYASWYQQKPGQAPVLVIYGKNNRPSGIPDRFSGSSSGNTASLTITGAQAEDEADYYCNSSPLNRLAVVFGGGTKLTVLG
-*SSSSGSSSSGSSSS*-GATTQQGSPVFAKLLAKNQASLCNTTLNWHSQDGAGSSYLSQGLRYEEDKKELVVDSPGLYYVFLELK
LSPTFTNTGHKVQGWVSLVLQAKPQVDDFDNLALTVELFPCSMENKLVDRSWSQLLLLKAGHRLSVGLRALHGAQDAYRD
WELSYPNTTSFGLFLVKPDNPWE

Supplementary Table 1: Sequences of the fusion proteins that were developed in this study

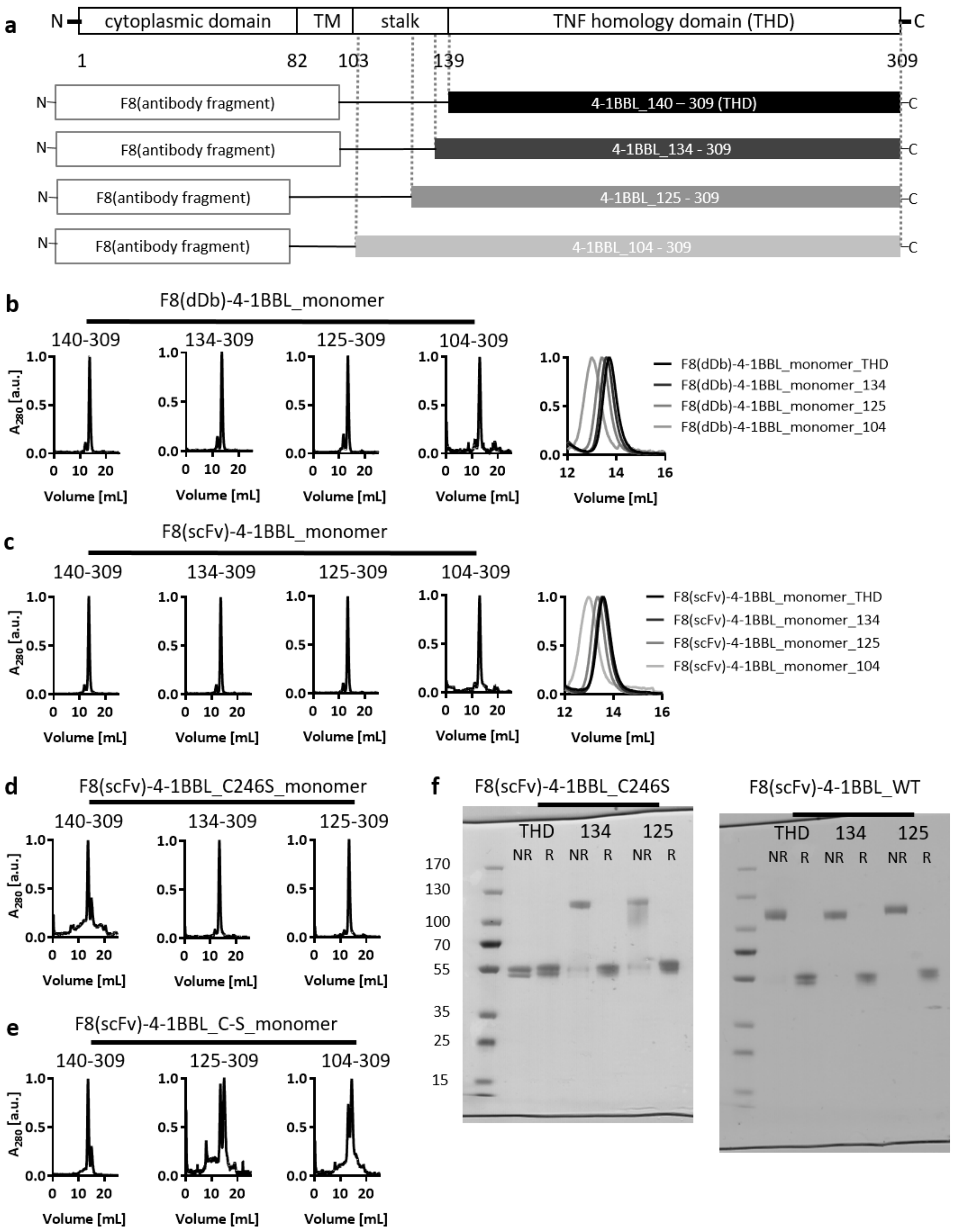


Supplementary Figure 1: Screening of the extracellular domain and C-S mutants of murine 4-1BBL to determine the optimal design of the 4-1BBL moiety in F8-4-1BBL (a) schematic depiction of the domain architecture of 4-1BBL and of the constructs that were screened containing different parts of the stalk region in addition to the TNF-homology domain (THD). The numbers correspond to the amino acid number in murine 4-1BBL. (b) Size exclusion profiles of F8 in the diabody (dDb) format linked to a single 4-1BBL subunit including different parts of the extracellular domain of 4-1BBL. Longer fragments of 4-

1BBL were more prone to aggregation and also gave lower expression yields (data not shown) (c) size exclusion profiles F8 in the scFv format linked to a single 4-1BBL subunit (d) Size exclusion profiles of different F8-4-1BBL variants featuring an C246S mutation in the 4-1BBL subunit preventing the formation of the disulfide bond that links two THD of 4-1BBL (e) Size exclusion profiles of different variants of F8-4-1BBL where all cysteines in the 4-1BBL domain were mutated to serines demonstrating the role of the disulfide bond for the stability of the fusion protein (f) SDS PAGE of the different C246S mutants revealing the presence of a second disulfide-forming cysteine outside the THD (NR: non-reducing, R: reducing)

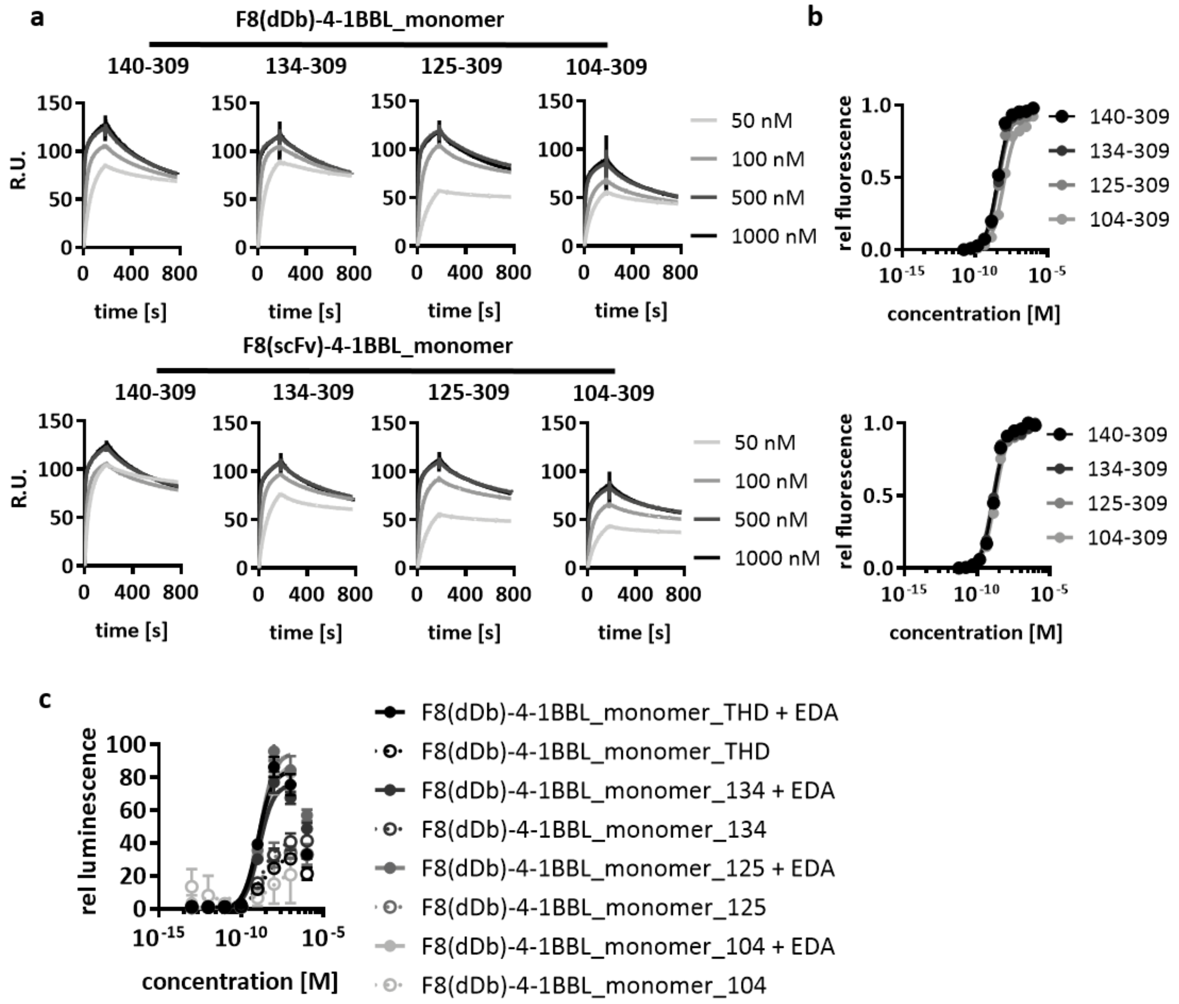


Supplementary Figure 2: The different F8-4-1BBL variants all retained binding to EDA and to 4-1BB as evidenced by (a) the Surface Plasmon Resonance measurements of binding to EDA and (b) the flow cytometry experiments with the 4-1BB expressing CTLL-2 cells (top: antibody in the diabody format, bottom: antibody in the scFv format) and (c) comparative bioactivity assays using the CTLL-2 NF-κB cell line revealed that the TNF homology domain is sufficient to achieve full signaling activity. The numbers refer to the amino acid numbers of murine 4-1BBL that were included in the F8-4-1BBL construct.

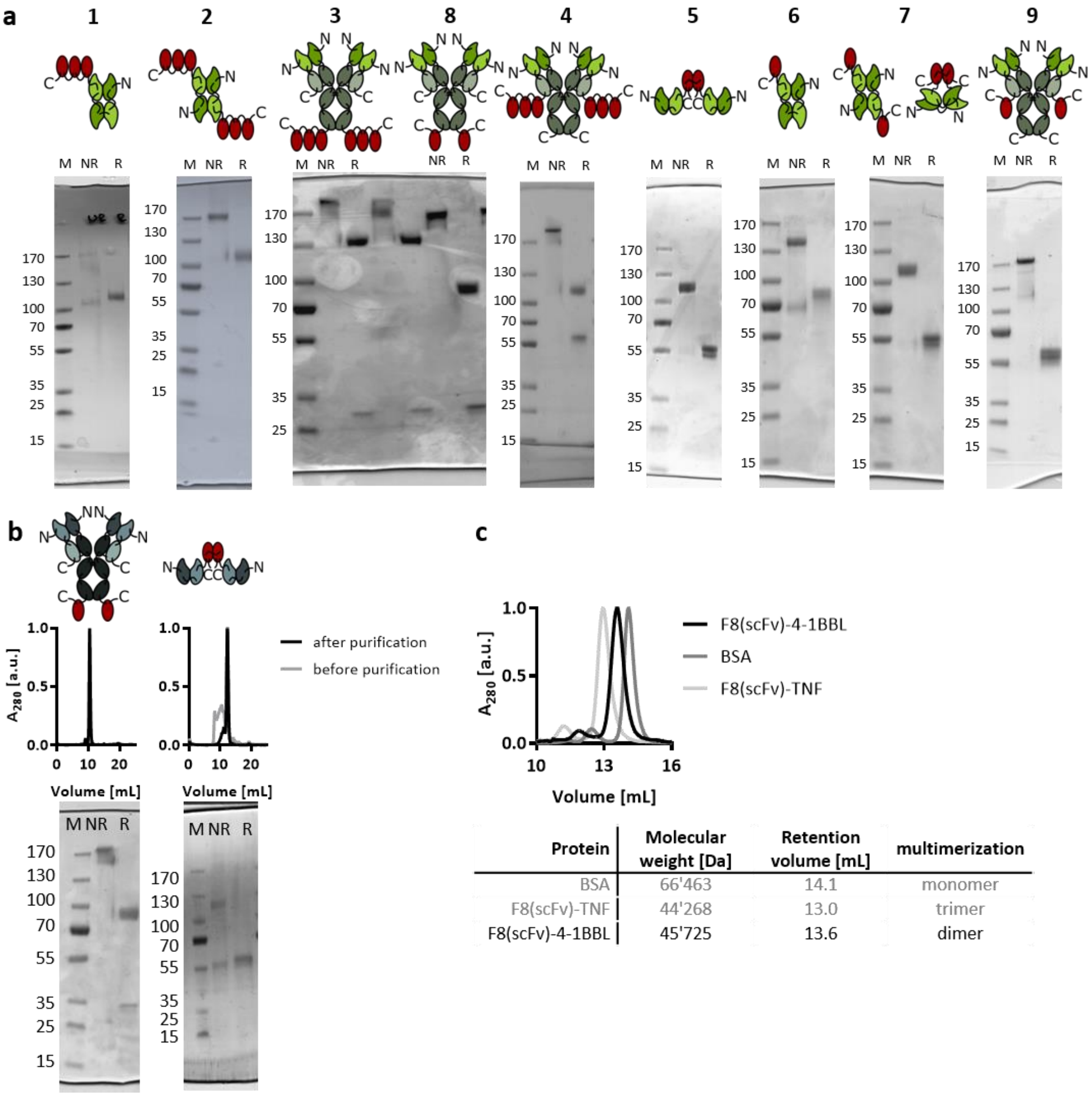


| Protein | Molecular weight [Da] | Retention volume [mL] | multimerization |
|---|---|---|---|
| BSA | 66'463 | 14.1 | monomer |
| F8(scFv)-TNF | 44'268 | 13.0 | trimer |
| F8(scFv)-4-1BBL | 45'725 | 13.6 | dimer |

Supplementary Figure 3: (a) SDS PAGE analysis of the nine F8-4-1BBL formats that were produced in this study (M: marker, NR: non-reducing sample buffer, R: reducing sample buffer) (b) Size exclusion chromatography profiles and SDS PAGE gels of the KSF constructs that were used for the biodistribution. The aggregates of the KSF(scFv)-4-1BBL could be efficiently removed by preparative size exclusion chromatography (c) comparative size exclusion chromatography showed that F8(scFv)-4-1BBL forms a dimer in solution since the retention volume was higher than for F8(scFv)-TNF that forms a non-covalent homotrimer[1]

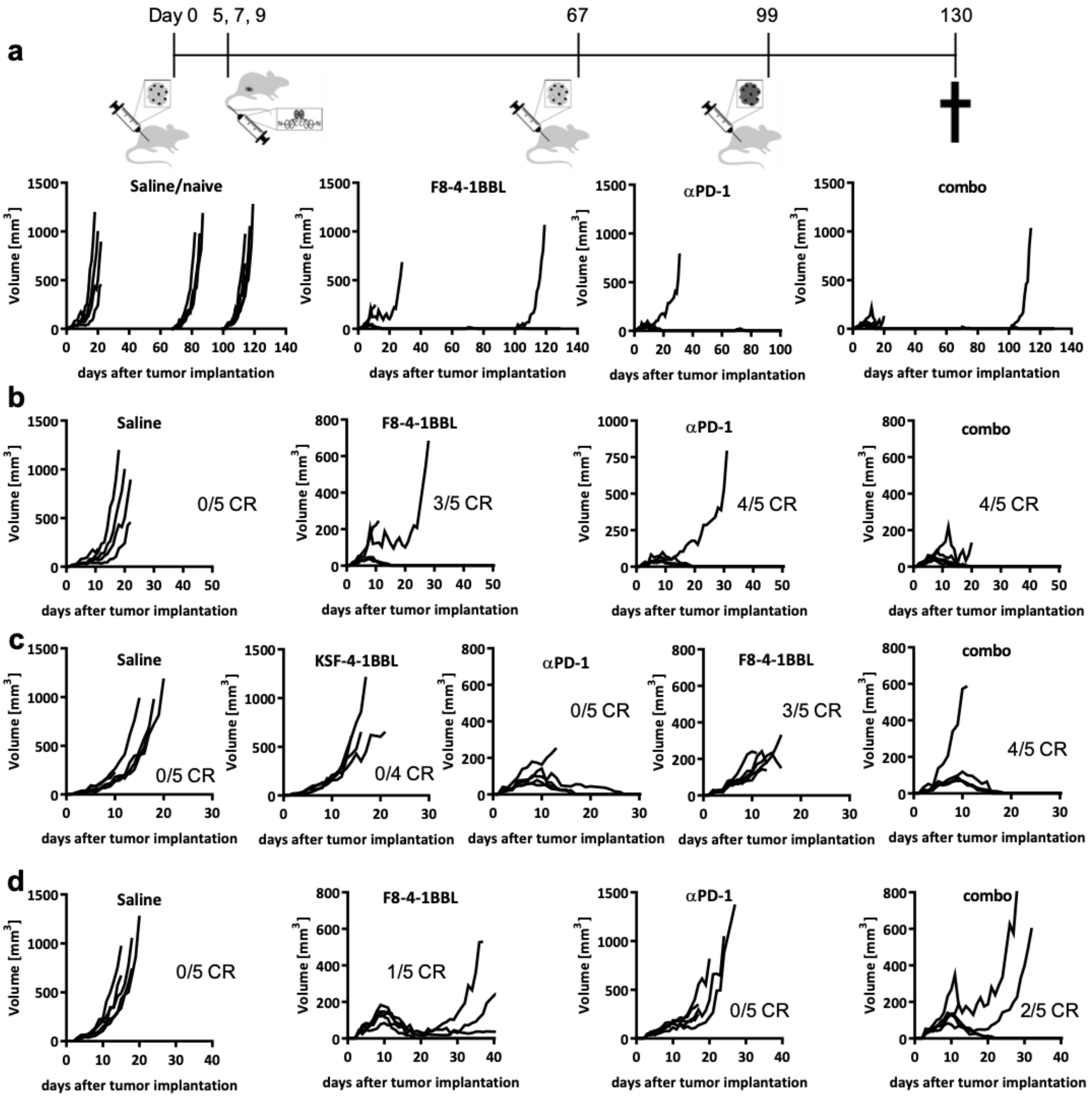


Supplementary Figure 4: Individual growth curves of the therapies (a) mice cured from WEHI-164 fibrosarcoma in the preventive setting were challenged with WEHI-164 fibrosarcoma cells on day 67 and mice from the F8-4-1BBL and the combo group also received a challenge with CT26 colon carcinoma cells on day 99. All the naïve mice developed tumors while the cured mice rejected subsequent challenges with WEHI-164 fibrosarcoma cells and most were also immune against CT26 colon carcinoma. (b) the individual growth curves of WEHI-164 fibrosarcoma bearing mice that received the treatment starting on day 5 after tumor implantation when the tumor volume was > 40 $mm^3$ (c) the individual growth curves of WEHI-164 fibrosarcoma-bearing mice that received the treatment starting on day 7 after tumor implantation when the tumor volume was > 80 $mm^3$ (c) individual growth curves of the CT26 colon-carcinoma-bearing mice that received the treatment starting on day 7 when the tumor volume was > 80 $mm^3$ (CR: complete response)

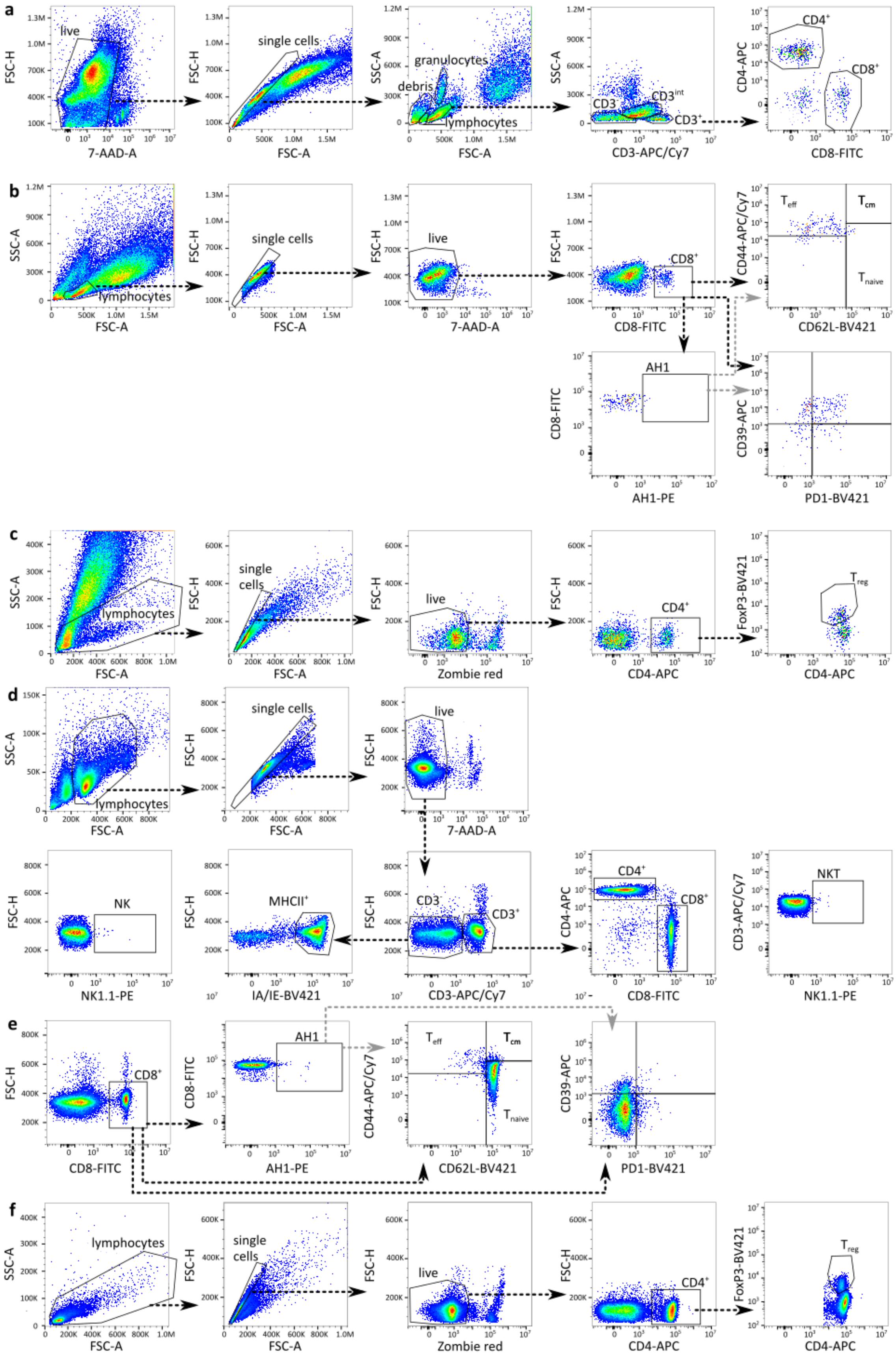


<u>Supplementary Figure 5:</u> Gating strategy for the flow cytometry analysis of the tumor-infiltrating lymphocytes (TILs) and the tumor-draining lymph nodes (TDLN) (a) in order to

quantify the tumor infiltrates, dead cells were excluded first by live/dead stain and then by scattering. Total living cells were calculated by subtracting dead cells and debris from the total number of events. Lymphocytes were further divided into the different subgroups (NKT, NK and MHCII+) as shown for the lymph nodes. (b) To assess the phenotype of tumor-infiltrating CD8+ cells, the events were gated by scattering and then live/dead, followed by CD8+ and AH1+. The same gates were applied to CD8+ and AH1+ cells for assessing the phenotypes. (c) To gate fixed tumor samples stained for intracellular FoxP3 the events were first gated by scattering and live/dead before the CD4+ cells were assessed for FoxP3 ($T_{reg}$). (d) To quantify the immune subsets in the tumor-draining lymph node (TDLN), the events were first gated by scattering and live/dead before the subsets were divided by CD3 expression and further markers. (e) The phenotype of CD8+ in the TDLN was assessed as described for the tumor. (f) Regulatory T cells in the TDLN were also assessed with the same gating strategy as described for the tumor (FSC: forward scattering, SSC: side scattering)

1 Hemmerle, T. *et al.* The antibody-based targeted delivery of TNF in combination with doxorubicin eradicates sarcomas in mice and confers protective immunity. *Br J Cancer* **109**, 1206-1213, doi:10.1038/bjc.2013.421 (2013).
